# Reliability of dynamic causal modelling of resting state magnetoencephalography

**DOI:** 10.1101/2023.10.16.562379

**Authors:** Amirhossein Jafarian, Melek Karadag Assem, Ece Kocagoncu, Juliette H Lanskey, Rebecca Williams, Yun-Ju Cheng, Andrew J Quinn, Jemma Pitt, Vanessa Raymont, Stephen Lowe, Krish D Singh, Mark Woolrich, Anna C Nobre, Richard N Henson, Karl J Friston, James B Rowe, the NTAD study group

## Abstract

This study assesses the reliability of resting-state dynamic causal modelling (DCM) of magneto-electroencephalography under conductance-based canonical microcircuit models, in terms of both posterior parameter estimates and model evidence. We use resting state magneto-electroencephalography (MEG) data from two sessions, acquired two weeks apart, from a cohort with high between-subject variance arising from Alzheimer’s disease. Our focus is not on the effect of disease, but on the predictive validity of the methods implicit in their reliability, which is crucial for future studies of disease progression and drug intervention. To assess the predictive validity of first-level DCMs, we compare model evidence associated with the covariance among subject-specific free energies (i.e., the ‘quality’ of the models) with vs. without interclass correlations. We then used parametric empirical Bayes (PEB) to investigate the predictive validity of DCM parameters at the between subject level. Specifically, we examined the evidence for or against parameter differences (i) within-subject, within-session, between-epochs; (ii) within-subject between-session and (iii) within-site between-subjects, accommodating the conditional dependency among parameter estimates. We show that for data acquired close in time, and under similar circumstances, more than 95% of inferred DCM parameters are unlikely to differ, speaking to mutual predictability over sessions. Using PEB, we show a reciprocal relationship between a conventional definition of ‘reliability’ and the conditional dependency among inferred model parameters. Our analyses confirm the predictive validity and reliability of the conductance-based DCMs for resting-state neurophysiological data. In this respect, the implicit generative modelling is suitable for interventional and longitudinal studies of neurological and psychiatric disorders.

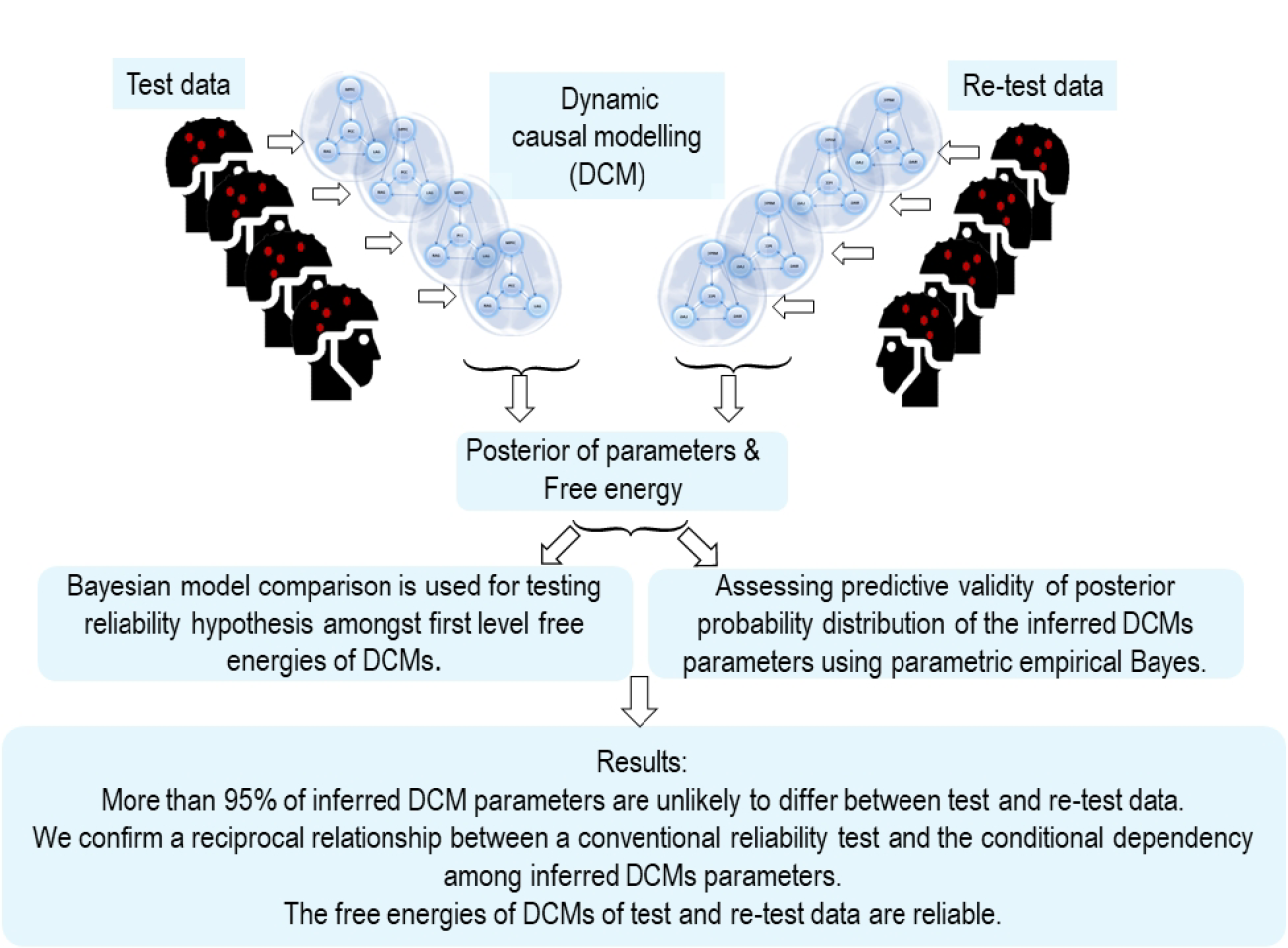

## 1 Introduction

Dynamic causal modelling (DCM) has been used widely in translational neuroscience to elucidate the underlying causes of physiological observations including electro/magneto-encephalography (MEG) (Adams et al., 2021a, Adams et al., 2022, Shaw et al., 2017, Shaw et al., 2021, Gilbert et al., 2016, Jafarian et al., 2023). However, the use of imaging and analytical methods to reveal the effects of disease progression and treatment intervention rests on reliability. This paper addresses the reliability of DCM, using variational Bayesian inversion of biologically informed models of neuroimaging data that can be used to characterise the neural mechanisms of cognition and the effect of disease or drugs. We test whether inferences from DCMs of resting-state MEG data are reliable i.e., predict each the results across trials within the same session, and across separate sessions from the same participants.

Dynamic causal modelling uses the variational Bayesian inversion of biologically motivated dynamical systems from neuroimaging data, to provide posterior estimates of unknown parameters (e.g. synaptic physiology) from a given model, and the model evidence (Friston et al., 2007, Friston et al., 2008). To compare alternative hypotheses, represented by alternative models, one uses differences in the free energy bound on [log-] model evidence, akin to log Bayes factors (Kass and Raftery, 1995, Friston, 2011, Friston et al., 2011, Jafarian et al., 2019). Bayesian model reduction (BMR) can be used for post hoc calculation of model evidence (and posterior parameter estimates) under nested, or alternative priors. Removing redundant parameters can improve model evidence by reducing model complexity. BMR is not only computationally efficient, but also eludes local minima during model inversion (Friston and Penny, 2011). At the group level, a hierarchal Bayesian inversion known as parametric empirical Bayesian (PEB) accommodates multiple first-level (single subject) models and constrains physiological parameters according to empirical priors quantifying between subject effects (Friston et al., 2015, Friston et al., 2016, Litvak et al., 2015). In this paper, we assess the reliability of these methods in DCM, in terms of predictive validity across trials and across sessions as measures of their reliability.

A classical statistical approach to measure reliability is the correlation between measures from the same participants under matched conditions (Fisher, 1992, Bartko, 1966). For two groups of data, (*x*_1_,*x*_2_)_*n*_ (*n* = 1,..*k*), the reliability can be defined as the modified Pearson correlation *r*, as follows :

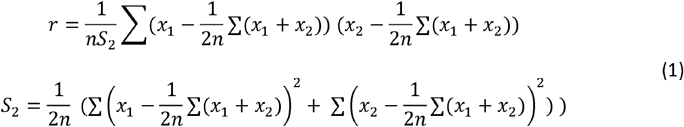

Conventionally, a general linear model or one-way analysis of variance (ANOVA) with random effects is used to quantify the reliability between measurements. If *y*_*ij*_ is the *i*^*th*^ measurements in the *j*^*th*^ group, ANOVA can be used to estimate the unknown group mean *μ*, unknown *j*^*th*^ group random effect *α*_*j*_ and normally distributed random effect ∈_*ij*_ in the following general linear model:

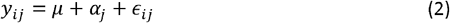

The between-group reliability is defined as 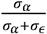, where the variance of group mean is *σ*_*α*_ and random effect variance is denoted by *σ*_∈_. However, as shown by (Box and Tiao, 1973, Wang and Sun, 2014), this point estimate of reliability is not always robust. Alternative reliability estimates using, for example, MCMC, can be limited by high computational burden (Mulder and Fox, 2019).

The frequentist approach to reliability considers only the expected values of parameters i.e. the maximum likelihood estimates, but not posterior variance or covariance. In ideal settings, frequentist estimates of DCM reliability may be sufficient. However, with complex models — with high posterior covariance among parameters — the reliability of the expected value of particular parameters can be very low (Schuyler et al., 2010, Frässle and Stephan, 2022, Rowe, 2010, Rowe et al., 2010). The reliability of DCM parameters has also been examined using split-sampling over evoked potentials (Adams et al., 2022) and resting state MEG (Jafarian et al., 2023). However, the frequentist approach to reliability is ill-suited to complex nonlinear dynamic systems with posterior covariance among model parameters. We therefore quantified reliability in terms of the contribution of subject or session effects using parametric empirical Bayes (PEB); effectively, comparing between subject or session models with and without between session or subject effects. This can be regarded as an assessment of predictive validity, in the sense that in the absence of between subject or session effects, the posterior estimates from one subject or session predicts the estimates from another.

This paper has three aims: (i) to use variational Bayes to test the reliability of evidence estimates (i.e., free energy) under DCMs inverted from MEG data, (ii) to leverage the PEB to test the reliability of inferred effective connectivity parameters; and (iii) test the influence of covariance among posterior parameters from DCM on the reliability of parameter estimates.

We use repeated measures of resting-state MEG data collected from participants in the “*New therapeutics in Alzheimer’s disease*’’ (NTAD) study (Lanskey et al., 2022). Our focus was not on modelling the effect of disease, but on the reliability of the ensuing estimates of synaptic efficacy. We briefly describe the participants and data, collected at baseline and two weeks later at rest. These data were acquired in a task-free or resting-state, which is suited for longitudinal studies of patients with progressive diseases. In the context of Alzheimer’s disease, we consider the oscillatory dynamics of the default mode network (comprising bilateral angular gyri, medial prefrontal cortex, and precuneus). Although medial temporal cortex is *a priori* associated with Alzheimer’s disease, MEG is relatively insensitive to activity in this region. Default mode network connectivity has been studied extensively in Alzheimer’s disease and its treatment (Lorenzi et al., 2011, Greicius et al., 2004). We performed first-level DCM to estimate synaptic parameters in the default mode regions, from the cross-spectral density of the MEG. We then consider the reliability of model evidence estimates, using a general linear model approach and the reliability of the DCM parameters using parametric empirical Bayes. Finally, we discuss the potential applications and limitations of the foregoing analyses. A glossary of acronyms and variables used in this paper is provided in the following tables.

## 2 Material and Methods

We tested the predictive validity or reliability of DCM estimates in terms of the evidence for between session (and subject) effects, inferred from repeated-measures data with conductance-based dynamic causal models.

### 2.1 Participants

The study received ethical approval from the Cambridge 2 Research Ethics Committee. Participants provided written informed consent. Participants met clinical diagnostic criteria for symptomatic Alzheimer’s disease, including mild cognitive impairment, with positive amyloid biomarkers (Lanskey et al., 2022). Principal participants undertook two MEG scans in a resting-state with eyes open, in two sessions that were two weeks apart — at a similar time of day — and without any medication changes. Their average age was 74.8 (standard deviation ±7.33), mini-mental state examination score was 26.1/30 (standard deviation ±3.05) and Addenbrookes Cognitive Examination (revised) score 78.5/100 (standard deviation ±10.5). A second set of participants underwent baseline MEG, with closely matched age 73 (±5.33) and cognition.

#### 2.1.1 Resting state MEG data

Five minutes of resting-state MEG data (with eyes open) were collected using an Elekta Vector View system with 204 planar gradiometers and 102 magnetometers. MEG data were recorded continuously with 1000 Hz sampling rate. Participants’ horizontal and vertical eye movements were recorded using bipolar electrooculogram and electro-cardiogram electrodes. Five head position indicator coils were placed on an EEG cap to track the head position. Three fiducial points (nasion, left and right pre-auricular) and >100 head shape points were digitised using Polhemus digitisation.

The Elekta Neuromag toolbox (Elekta Oy), with MaxFilter v2.2.12 was used for the detection and interpolation of bad sensors, signal space separation to remove external noise from the data and head movement correction. Then we pre-processed the data by down-sampling to 500 Hz, bandpass filtering between 0.1-100 Hz, and applying a notch filter between 48-52 Hz and 98-102 Hz. We then performed artefact rejection/removal using ICA, with EOG data. We epoched the data into 1000 ms segments and repeated each epoch’s artefact rejection and removal.

Using T1-weighted structural MRI (3T Siemens, TR = 2300 ms, TE = 2.91 ms, resolution 1 mm), we performed DICOM conversion to NII and inverse-normalised the canonical mesh, size 2. We co-registered the MRI to the mesh using three fiducials and head shape points to create the forward model for MEG with the single shell boundary element model method. We used “COH” source inversion (Litvak et al., 2011) for extracting four default mode network sources/regions in left and right angular gyri (LAG [49 -63 33], RAG [-46 -66 30]), medial prefrontal cortex (MPFC) [-1 54 27], and Precuneus (PPC) [0 -55 32]. We used the induced source inversion option over 1000ms epochs, frequency range 0.1-100 Hz, with fusion across magnetometers and gradiometers. We extracted principal components of power spectral densities over trials, as data features for the DCM of cross-spectral density.

We use three sets of resting-state eyes open MEG data for the reliability study: (i) split sampled baseline data where each individual patients is divided into odd and even epochs, (ii) baseline vs two weeks data, and (iii) data from baseline and a second set of participants at baseline.

### 2.2 Dynamic causal modelling of resting-states MEG data

We use DCM for cross-spectral density (CSD), SPM12-version 8163 (Friston et al., 2012, Moran et al., 2007, Moran et al., 2011) for inferring parameters and the marginal likelihood of the conductance-based biophysically canonical microcircuit model (*cmm-nmda model in SPM12*) from spectral features of MEG data. DCM for CSD explains the frequency content of MEG data in terms of a local linear perturbation (due to endogenous neuronal fluctuations) around the fixed point of a nonlinear model of canonical neuronal circuitry (Basar et al., 2012, Haken, 1977).

The conductance-based model describes the electrical activity of a cortical source based on the interactions of four neuronal populations: excitatory spiny stellate cells, superficial pyramidal cells, inhibitory interneurons, and deep pyramidal cells, as shown in Figure 1. Each cortical source is connected to other regions via forward connections (that originate from the superficial pyramidal population and project to excitatory spiny stellate and deep pyramidal cells of other regions) and backward connections (that originate from deep pyramidal cells and project to superficial pyramidal and inhibitory populations in the target source). Each population is modelled by a Morris–Lecar model (driven by random endogenous fluctuations) (Moran et al., 2013) as follows:

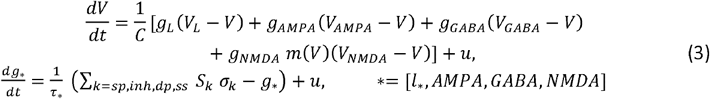

**Figure 1.**
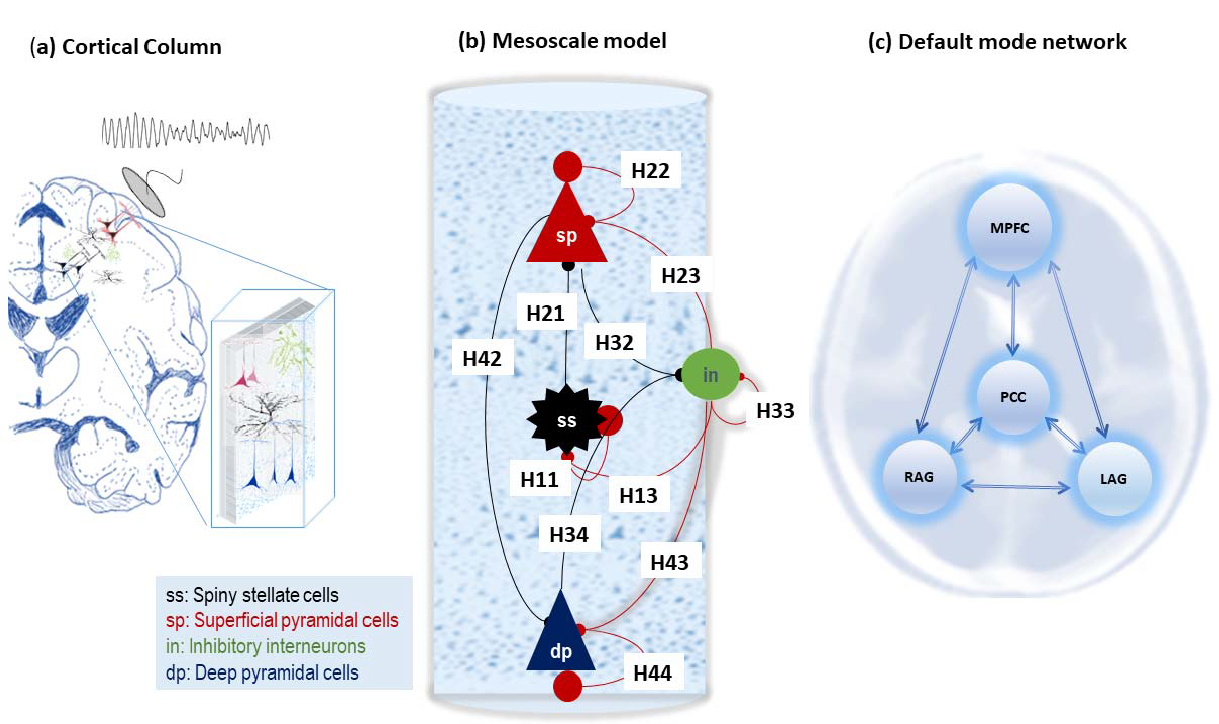
Mesoscale model of MEG data. Panel (a) illustrate a cortical column as the origin of electrical brain activity as recorded by neuroimaging modalities such as MEG. Panel (b) illustrates the laminar specific conductance based model with superficial (sp) and deep pyramidal (dp) cells in the top and bottom layers, respectively, excitatory interneurons (spiny stellate cells, ss) situated in layer four, and inhibitory interneurons that are distributed across all layers and modelled using one population. The dynamics of ion compartments with a population are governed by the Morris-Lecar model. This model explains the dynamics of different ion currents: NMDA, AMAP and GABA and passive ion current and membrane capacitance as explained in equation 1. Panel (c) shows the fully connected default mode network which contains MPFC. PCC, RAG and LAG sources.

In Equation 3, the membrane potential is denoted by *V*, the conductance of an ion channel is *g*_*L,NMDA,AMPA,GABA*_, and *u* is random endogenous fluctuation. Constant parameters are *C* as the membrane capacitance, *l*_*_ as a passive leak current with a fixed conductance, and *τ*_*_ as ion channel receptor time constants. *V*_*L,NMDA,AMPA,GABA*_ denote the reversal equilibrium potentials of the ion channels. The dynamics of depolarisation is equipped with activity-dependent magnesium channels, which are modelled as 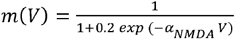 Afferent presynaptic firings from a population *k* are denoted by *σ*_*k*_, which is scaled by connectivity *S*_*k*_ (which can be, in nature, excitatory AMPA, NMDA or inhibitory GABA afferent intrinsic *H*_*k*_ or forward and backward extrinsic AMPA (denoted by A) and NMDA (denoted by *AN*) connections.

The generative model of MEG data (denoted by *y*) can be written as a partially observed dynamical system as follows:

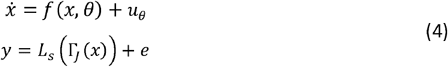

In equation 4, *θ* denotes a lump representation of all unknown parameters (i.e., *H*_*k*_, *τ*.), *x* is a vector of biological states in the model, *f*(*x, θ*) is a function that is the concatenated version of the right-hand sides of Equation 1 over all populations. In Equation 4, *u* is the endogenous (structured pink) noise with a cross-spectrum, *g*_*u*_(*ω, θ*) = *FT*(*E*[*u*(*t*), *u*(*t* − *τ*)]). In the second line of Equation 4, *L*_*S*_ is a parameterisation of lead field projections and sensor gain, and *Γ*_*J*_(*x*) is an operator (parameterised by unknown parameters *J*) that links the activity of populations (e.g., weighted sum — by *J* parameters — of different population responses) to MEG data (through lead field projection).

Resting state dynamics can be modelled as a response to endogenous random fluctuations. In this DCM, a linearised neuronal model (around a stable equilibrium point) is used to generate spectral content of the MEG data. The spectral response of the neuronal model, *g*_*x*_(*ω*), can be modelled as follows:

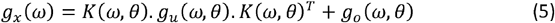

In Equation 5, *g*_*o*_ (*ω, θ*) represents the spectrum of the observation noise, which is a sum of common and source-specific noise and *K* (*ω, θ*) = *FT*(exp *τ*·*∇*_*x*_*f*(*x, θ*)) (*∇*_*x*_ is the Jacobian) is the transfer function (input to output) of the neural model (parametrised by *J*). The spectral response in sensor space can also be generated by the inclusion of a forward electromagnetic model into Equation 4, which is denoted by *L*.*M* (*L* is the sensor gain and *M* is the head model), as follows:

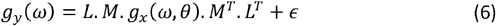

In Equation 6, *g*_*o*_ (*ω*) is the cross spectra of the MEG data, and *∈* ∼ *N*(0, *σ*^2^) is a random effect (with unknown covariance). Because we perform the DCM in the source space, the forward electromagnetic model reduces to a scaling parameter.

The laminar interpretation of the conductance-based model (e.g. superficial, deep geometry) is supported by (i) specification of priors for intrinsic connections (Table 1), (ii) the equation of the observer for MEG data and (iii) endogenous inputs to the model which targets spiny stellate cells in layer four. This model parametrisation supports inferences about the laminar basis of degenerative brain neurological disorders (Adams et al., 2021a, Adams et al., 2022, Adams et al., 2021b, Shaw et al., 2021).

**Table 1:**
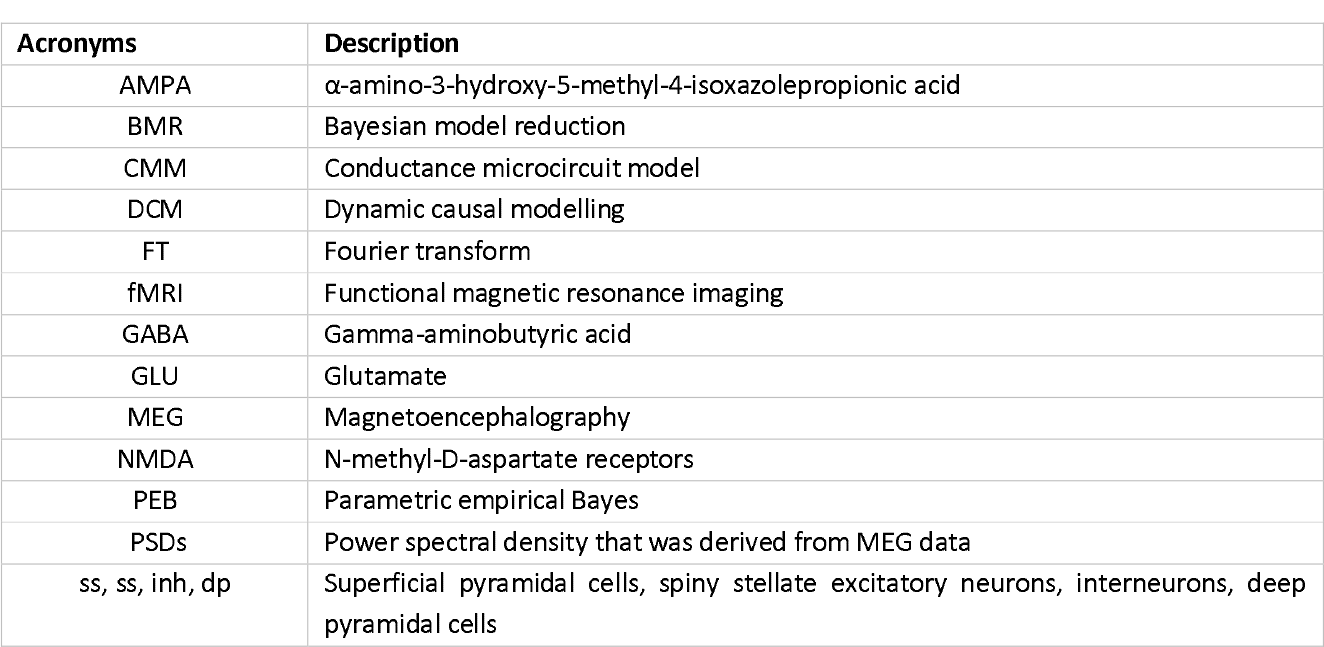
Acronyms.

| Acronyms | Description |
| --- | --- |
| AMPA | α-amino-3-hydroxy-5-methyl-4-isoxazolepropionic acid |
| BMR | Bayesian model reduction |
| CMM | Conductance microcircuit model |
| DCM | Dynamic causal modelling |
| FT | Fourier transform |
| fMRI | Functional magnetic resonance imaging |
| GABA | Gamma-aminobutyric acid |
| GLU | Glutamate |
| MEG | Magnetoencephalography |
| NMDA | N-methyl-D-aspartate receptors |
| PEB | Parametric empirical Bayes |
| PSDs | Power spectral density that was derived from MEG data |
| ss, ss, inh, dp | Superficial pyramidal cells, spiny stellate excitatory neurons, interneurons, deep pyramidal cells |

**Table 2:** Glossary of variables and expressions in the conductance-based model (CMM-NMDA)

| Variable | Description |
| --- | --- |
| $u$ | Exogenous input |
| $V$ | Mean depolarisation of a neuronal population |
| $\sigma(v)$ | The neuronal firing rate – a sigmoid squashing function of depolarisation |
| $L$ | Lead field vector mapping from (neuronal) states to measured (electrophysiological) responses |
| $g_x(\omega), g_o(\omega), g_y(\omega)$ | Spectral density of (neuronal) state fluctuations, observation noise and measurement, respectively |
| $\nabla_x f$ | System Jacobian or derivative of system flow with respect to (neuronal) states |
| $k(t) = FT[K(\omega)]$ | First-order kernel mapping from inputs to responses; c.f., an impulse response function of time. This is the Fourier transform of the transfer function |
| $K(\omega) = FT[k(t)]$ | The frequency transfer function modulates the power of endogenous neuronal fluctuations to produce a (cross-spectral density) response. This is the Fourier transform of the kernel |

**Table 3:** Parameters of the neuronal model (see also Figure 1). The (*i, j*) element in the matrix associated with parametrisation of intrinsic connection *H*, means connections that originate from population *j* and target population i in a region (here element 1 to 4 are corresponds to ss, sp, inh and dp layers respectively). The (*i, j*) element in the matrix associated with parametrisation of extrinsic connections *A* and *AN*, means connections that originate from population *j* in a region and project to population *i* in a distal region (here element 1 to 4 are corresponds to ss, sp, inh and dp layers respectively).

|  | Description | Parameterisation | Prior |
| --- | --- | --- | --- |
| $\kappa$ | Rate constants of ion channels, AMPA, GABA, and NMDA respectively | $\exp(\theta_\kappa) \cdot \kappa$<br>$\kappa = [4, 16, 100]$ | $p(\theta_\kappa) = N(0, 1/16)$ |
| $C$ | Membrane capacitance of ss, sp, inh, and dp populations respectively | $\exp(\theta_c) \cdot C$<br>$C = [128 \ 128 \ 256 \ 32] / 1000$ | $p(\theta_c) = N(0, 1/16)$ |
| $H$ | Intrinsic connections | $\exp(\theta_H) \cdot H$<br>$H = \begin{bmatrix} 8 & 0 & 2 & 0 \\ 4 & 8 & 8 & 0 \\ 0 & 0 & 32 & 128 \end{bmatrix}$ | $p(\theta_H) = N(0, 1/32)$ |
| $A$ | Extrinsic forward connection | $\exp(\theta_A) \cdot A$<br>$A = [1 \ 0; 0 \ 1; 0 \ 2; 0 \ 0]/8;$ | $p(\theta_A) = N(0, 1/8)$ |
| $AN$ | Extrinsic backward connection | $\exp(\theta_{AN}) \cdot AN$<br>$AN = [1 \ 0; 0 \ 1; 0 \ 2; 0 \ 0]/8;$ | $p(\theta_A) = N(0, 1/8)$ |
| $L$ | Sensor gain | $L$ | $p(L) = N(1, 64)$ |
| $J$ | Contribution of spiny stellate population ($J_{ss}$) and deep pyramidal ($J_{dp}$) to observation data | $J_*, (* = dp, ss)$ | $p(J_{ss}) = N(0, 1/16)$<br>$p(J_{dp}) = N(0, 1/16)$ |
| $a$ | Endogenous random fluctuation with transfer function $\frac{a_1}{\omega^{a_2}}$ | $\exp(a)$ | $p(a_{1,2}) = N(0, 1/128)$ |
| $d$ | Structural cosine coefficients of endogenous random fluctuation | $\exp(d)$ | $p(d_{1,2,3,4}) = N(0, 1/128)$ |
| $b$ | Common sensor noise with transfer function $\frac{b_1}{\omega^{b_2}}$ | $\exp(b)$ | $p(b_{1,2}) = N(0, 1/128)$ |
| $c$ | Specific sensor noise with transfer function $\frac{c_1}{\omega^{c_2}}$ | $\exp(c)$ | $p(c_{1,2}) = N(0, 1/128)$ |
| $f$ | Scaling some frequencies as model of data filtration | $\exp(f)$ | $p(f_{1,2}) = N(0, 1/128)$ |
| $D$ | Neuronal delay between regions and within layers | $\exp(D) \cdot D$<br>$D = [2, 16]$ | $p(D) = N(0, 1/64)$ |

The unknown parameters in the DCM are specified as log-scale values 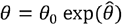 where *θ*_*o*_ are biologically informed scaling constants for the parameter, and 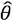 has Gaussian distribution, with prior normal density 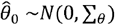 of zero mean and covariance ∑_*θ*_. This expresses Bayesian beliefs about the range over which parameters can vary and constrains the posterior density over parameters accordingly. The prior distribution for parameters assures the stability and plausibility of the model, and we refer to them as “micro-priors” hereafter.

### 2.3 Parametric empirical Bayes

Dynamic causal modelling at the between session or subject level implements hierarchical variational Bayesian inversion of data under empirical priors at the second level (e.g. age, disease severity, etc) on the first level (e.g., synaptic) parameters (please see Appendix A for a mathematical treatment of parametric empirical Bayes or PEB). At the first level, the neuronal model is optimised to fit each individual’s data. At the second level, the estimation of some parameters is constrained by group estimates to improve group model evidence. These parameters are those that are equipped with random effects. For these parameters there are micro- and macro-priors that constrain the accompanying posterior estimates at the first and second level, respectively.

The micro priors are taken from the physiological literature (Friston and Penny, 2011, Friston et al., 2015, Friston et al., 2003), while macro priors impose constraints on parameters based on information about the population from which the data are drawn (i.e. an empirical prior, such as age or disease severity) (Friston et al., 2016). Due to leveraging Bayesian model reduction, assessing the impact of macro-scale constraints on synaptic parameters does not require re-estimation of the first level DCMs (Friston et al., 2016, Friston et al., 2019, Zeidman et al., 2019).

### 2.4 Reliability of dynamic causal modelling

#### 2.4.1 Reliability of fixed quantities

This section assesses the reliability between fixed effects (i.e., measurements without uncertainty or random effects), such as the free energy of a model. We denote two sets of measurements of similar phenomena by column vectors *θ*_1_ and *θ*_2_, each of which has dimension *n* × 1 (*n* number of, e.g., participants) and specify the following linear model:

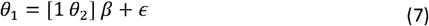

In Equation 7, *β* represents the intercepts of the model between the two measurements. The reliability between measurements can be specified as a covariance component of the noise term *∈* ∼ *N*(0, Σ), with compound symmetry as follows (Jelenkowska, 1998, Jelenkowska, 1999)

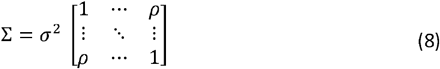

The covariance matrix elements are defined as [Σ_*ii*_] = *σ*^2^ and [Σ_*ij*_] = *ρσ*^2^(*i* ≠ *j*). The parameter *ρ* is the inter-class correlation coefficient.

To test reliability, one can invert the general linear model in Equation 7 with and without *ρ* in the covariance matrix, and use Bayesian model comparison to compare the respective free energies to assess the evidence for an interclass correlation. We use variational Laplace (Friston et al., 2007) as implemented in SPM12 to estimate the covariance components, with and without the interclass correlation.

#### 2.4.2 Reliability of inferred DCM parameters

To assess the reliability of parameters with random effects, we used a parametric empirical Bayesian approach to assess the contribution of random effects to condtionally dependent parameter estimates.

Motivated by the classical definition of reliability, i.e. that there are no between-session or between-subject effects, we evaluated the evidence for PEB models with and without these effects. We perform PEB estimation for each parameter separately; in which synaptic rate constants (*T*), intrinsic synaptic gain (*H*), extrinsic connections (*A* and *AN*), state to observation parameters (*L* and *J*), and physiological inputs (*a* and *b*) are constrained by their group average (a single column matrix of ones). We then used the PEB of PEB approach, with the following design matrix to assess the evidence for between session or subject effects:

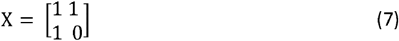

## 3 Results

The power spectra of the MEG data from each source and subject are shown in the supplementary material. Comparison between the baseline and two-week spectra show that they are not identical. These differences are not attributable to the progression of the disease over such a short interval but may rise from other factors, including for example psychological states (e.g. fatigue), recording noise, or movements.

### 3.1 Reliability of DCM

We tested the reliability of the model’s free energy estimates based on test and re-test data: as split-sample data from one session, or two sessions acquired in the same participants, or of two similar groups of participants, as summarised in Figure 2. The free energy estimates were highly reliable for within-session split-sample data, and within-subject between session data. For the within-subject models, the free energy was higher for the model with compound symmetry — i.e., with a high interclass correlation — with a difference of 20 (equivalent to a Bayes Factor of ∼5×10^8^). On the other hand, there was no evidence for an interclass correlation between the free energy of models of data acquired from different people, even if demographically and clinically matched. This is expected: even though most of the neuronal parameters of the biophysical models that generate MEG data were similar for matched adults, differences in signal-to-noise and other non-neuronal factors can have a profound effect on the free energy estimates of model evidence (a.k.a., marginal likelihood).

**Figure 2.**
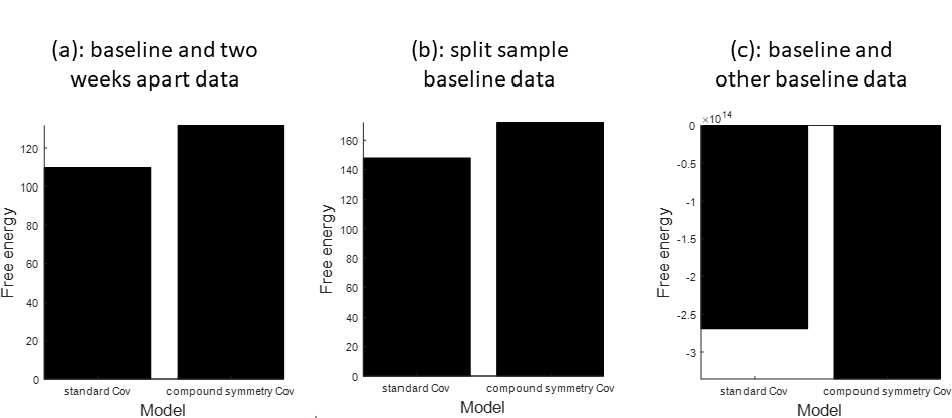
Reliability of free energies of the first level multichannel DCMs of the default mode network. There is greater evidence for the model with compound symmetry structure than the model without, when fitting data for different trials within-subjects (panel a) or different sessions within-subjects (panel b; Baseline vs Two-week data), but greater evidence for the model without compound symmetry when fitting data from different subjects, as expected (panel c; Baseline data).

To test for between session and subject effects, we used PEB of PEBs to assess the reliability of 156 physiological parameters; including synaptic rate constants (*T*), intrinsic synaptic gains (*H*), extrinsic forward and backward AMPA connections (denoted by *A*) and extrinsic forward and backward NMDA connections (denoted by *AN*), state to observation parameters (*L* and *J*), and physiological inputs (*a* and *d*). For the within-session split-sample analysis, the PEB of PEBs revealed that only four of 156 parameters differed between odd and even trials (Figure 3-a). For the within-subject between-session analysis, the PEB of PEBs showed that only four of 156 parameters differed between sessions (Figure 3-b). Finally, there was fair agreement across the two separate groups of similar participants, with nine of 156 showing evidence of between subject effects on parameter estimates (Figure 3-c).

**Figure 3.**
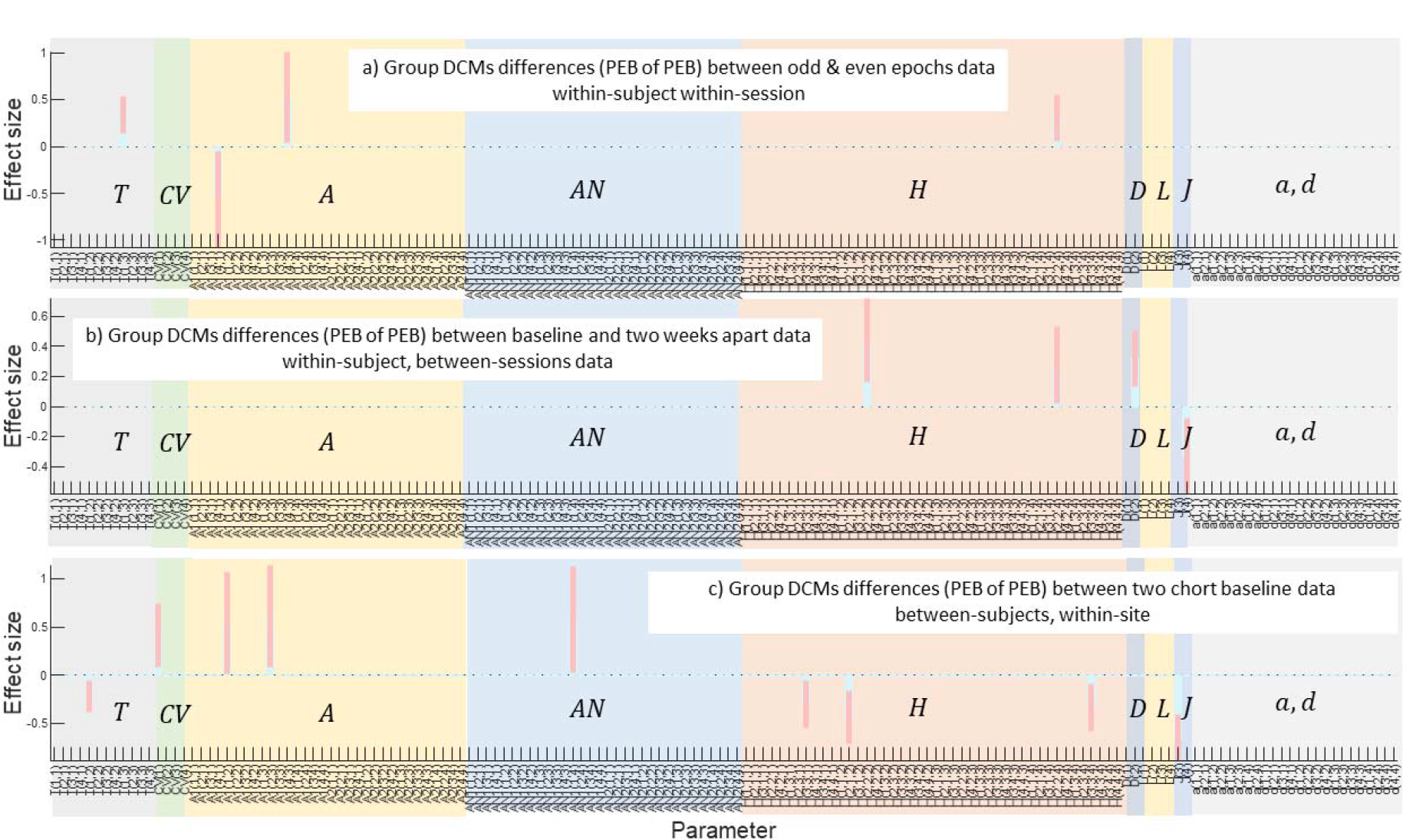
Reliability using the PEB of PEB approach for 156 inferred parameters of the fully connected default model network DCMs for (a) split sample data within-subject within-session, (b) within-subject between-sessions, two weeks apart and (c) between-subjects, within-site. Each plot illustrates the expectation of parameters (blue bar) and their 95% confidence interval (pink bar) for parameters for which there was evidence of a difference. Note that >95% of parameters do not differ within-session, or between-sessions. Time constants are denoted by T and CV are membrane capacitances, A and AN are between-sources forward and backward extrinsic AMPA and NMDA connections, respectively, H are the intrinsic within-regions connections, are within and between regions delays, L and J are sensor gains and states that contributed to local field potentials respectively and a and b are parameters of random neuronal fluctuations.

### 3.2 Reliability and the correlation of parameters

To quantify the relationship between the reliability of the posterior parameters estimates and their correlation, we examined the expected values of parameters after first level DCM and their re-estimation by the PEB, i.e., second level DCM to implement empirical priors or constraints. In Table 4, we report number of intrinsic, extrinsic forward and backward synaptic parameters that have correlations larger than 0.5 between their first-level DCM estimates and after re-estimation by PEB under empirical (second level) constraints. This confirms that after implementing empirical priors, their reliability improves. The PEB re-estimates first-level DCM parameters to improve model evidence by reducing model complexity and implicitly resolving conditional dependencies. In effect, inferred parameters become more conditionally independent, and thereby more reliable in the classical sense.

**Table 4:** Reliability of parameters before and after using PEB for split sampled and follow up data.

| Inferred DCM parameters | Number of parameters with Correlation score > 0.5 in DCMs of split sample baseline data |  | Number of parameters with Correlation score > 0.5 in DCMs of two weeks apart data and baseline data |  |
| --- | --- | --- | --- | --- |
|  | First level DCM | Re estimated first level DCM by PEB | First level DCM | Re estimated first level DCM by PEB |
| Intrinsic connections H | 15 | 19 | 8 | 10 |
| Extrinsic forward connection A | 8 | 9 | 7 | 8 |
| Intrinsic backward connections AN | 9 | 10 | 11 | 12 |

## 4 Discussion

We assessed the reliability of dynamic causal modelling using magnetoencephalography data acquired within-session, between-session and between-subject. In all three cases, the conductance-based DCM estimates, obtained from resting state MEG power spectra, are highly reliable in terms of the inferred neuronal parameters. To compare fixed quantities (e.g. the free energy estimates of model evidence), we used variational Laplace to estimate the covariance components due to intra-class correlations. To assess agreement between parameters with random effects, we used “PEB of PEBs” to test for between-session and between-subject effects, finding excellent generalisation over split-sample and test-retest analyses. These indicate a high reliability of DCM of electrophysiological observations. In addition, we confirmed the reciprocal relationship between reliability and conditional dependency (covariance) between parameter estimates. In other words, as the covariance among parameters reduces, the reliability of their expectations increases.

A key motivation for this analysis was to address the reliability of inferences from complex biophysical models in which neuronal generators are nonlinear, and parameter estimates are correlated (Litvak et al., 2019). In previous studies, individual DCM connectivity parameters have proven unreliable, except for very simple models, despite the high reliability of model evidence and model comparison for hypothesis testing (Schuyler et al., 2010, Jafarian et al., 2022, Rowe, 2010, Rowe et al., 2010). Poor reliability was attributed to their covariance, and the use of model comparison based on model evidence was recommended for hypothesis testing.

Parameter inference is sensitive to changes in the data (which may arise in repeated-measures or between-subject data). However, using PEB (Friston and Penny, 2011, Friston et al., 2016, Litvak et al., 2015, Litvak et al., 2019) one can effectively identify the similarity of underlying neuronal dynamics between split-sample and test-re-test data for ∼95% of parameters; consistent with the data being drawn from the same distribution, and in our study the absence of disease progression.

Parameter estimates under nonlinear models are accompanied by a degree of co-linearity that is, changing one parameter is equivalent to changing others. In other words, different combinations of parameters in the generative model could give rise to very similar data. The set of parameters can be considered as a manifold, which is biologically plausible (Prinz et al., 2004) but challenging when testing hypotheses based on particular parameter values. Historically, solutions to this problem have included reducing model complexity (Stephan et al., 2007), by re-parameterisation and defining priors for the generative models for functional MRI. However, such re-parameterisation is not straightforward, and may not be possible for generative models of M/EEG. This is because of the inherent complexity of the models of M/EEG cortical generators (Penny, 2012). To address this problem, we leveraged BMR and PEB to find nested models with lower complexity that can better capture the underlying dynamics.

The use of PEB can be seen as an extension of BMR to cohort studies. PEB is well suited to address whether model evidence from cohort data can be improved by replacing non-informative (or weakly informative) prior parameters in DCM by empirical priors; i.e., empirical constraints (Jafarian et al., 2023, Jafarian et al., 2021, Adams et al., 2022, Adams et al., 2021b, Friston et al., 2022). In PEB, a reduced model with higher model evidence is sought at the group level by constraining parameters using group information to enhance cohort model evidence. PEB re-estimates first-level DCMs where all parameters are informed by the group information. If two sets of data are sampled from the same distribution, as expected within-session, then PEB of PEB should not identify differences in the parameters of the generative model. The improvement of reliability after the application of PEB is partly due to the application of Bayesian model reduction during group DCM inversion, which serves to reduce the complexity and, implicitly, the posterior correlation between parameters.

The reliability of the inferred parameters and model evidence from DCMs is just one aspect of their validation. It is not expected that the inferred parameters are identical, given the likely differences between the data acquired close together and in ‘similar’ conditions. Although disease progression is unlikely to a meaningful degree in two weeks, other differences like fatigue, anxiety, motion or scanner noise, may arise. These differences are likely to occur in addition to longitudinal or interventional “repeated-measures” studies. The effect of such cofounds for translational neuroscience is a matter of degree, and approaches to reduce the effect of such confounds include (i) using larger data samples and (ii) inclusion of informative priors on individual differences (Stephan et al., 2009, Jafarian et al., 2023, Zeidman et al., 2019).

In summary, we have demonstrated reliable estimates of inferred parameters and relative model evidence, both within-session and between-sessions. The use of group inversion improves the free energy of first-level DCMs and improves reliability of individual parameters. Such DCMs, including the canonical microcircuit model of magnetoencephalography, provide a sufficiently reliable modelling platform to consider for use in longitudinal or interventional studies.

### 5 Appendix

#### 5.1 Mathematical treatment of PEB

Mathematically, let a column vector of model parameters at the first level DCM, over cohort, is 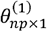 (*n* number of participants and *p* is the number of parameters for each participant). Then the generative model of the PEB is given by (Friston et al., 2016):

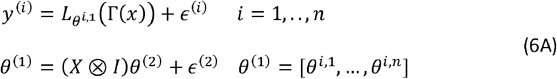

The first line of equation 6A is the generative model of each DCM with unknown parameters inferred from neuroimaging data at the first level. The second line models (macro-level) empirical priors that constrain parameter estimates from the first-level DCM. In the second line of equation 6A, *X* ∈ *R*^*n*×*r*^ is the design matrix with *r* ≥ 1 covariant. The first column of *X* is equal to one and reflects the group mean; generally, the rest of the column can be defined based on empirical data. The symbol ⊗ is the Kronecker product, and *I* is the *p* × *p* identity matrix. The random effects have a Gaussian distribution and at the second line of equation (6A), *∈*^(2)^ ∼ *N*(0, *Π* ^(2)^) (where *Π*^(2)^ is the precision matrix or inverse of covariance). The precision matrix is parameterised with a single (hyper-precision) parameter, *γ*, as follows (Friston et al., 2016):

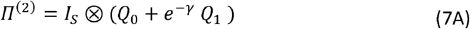

In equation 7A, *Q*_0_ ∈ *R*^*p*×*p*^ is the lower bound on the precision, defined with a small positive value. The (hyper)parameter, *γ*, scales a precision matrix *Q*_1_ ∈ *R*^*p*×*p*^, which is (by default) 16 times the prior precision of the group mean (Zeidman et al., 2019): this hyper prior ensures that random effects are small compared to prior uncertainty about the parameter in question.

The objective of group DCMs is to maximise the second level free energy under the PEB constraint. The inversion of group DCM starts with the inversion of each data, followed by assessing the effect of empirical prior and adjusting parameters so that the free energy is maximised. As part of group DCMs inversion, BMR is used to re-evaluate first-level posteriors under updated second-level parameters (Friston et al., 2016, Litvak et al., 2015). This greatly expedites inversion of group data.

## 6 Contribution

**Amirhossein Jafarian:** Conceptualization, Methodology, Software, Validation, Formal analysis, Writing Original Draft, Data Curation, Visualization. **Melek Karadag Assem:** Data acquisition, Pre-processing of MEG data, Review & Editing. **Ece Kocagoncu:** Data acquisition, Pre-processing of MEG data, Review & Editing. **Juliette H Lanskey:** Data acquisition, Review & Editing, **Rebecca Williams:** Review & Editing, **Andrew J Quinn:** Data acquisition, Review & Editing, **Jemma Pitt:** Data acquisition, Review & Editing, **Stephen Lowe:** Review & Editing, **Vanessa Raymont:** Review & Editing, **Krish D Singh:** Review & Editing, **Mark Woolrich:** Review & Editing, **Anna C Nobre:** Discussion, Review & Editing, **Richard N Henson:** Discussion, Review & Editing, **Karl J. Friston:** Conceptualization, Methodology, Review & Editing. **James B Rowe:** Conceptualization, Methodology, Review & Editing, Funding acquisition.

## 7 Acknowledgements

This work has been funded by the Wellcome Trust (220258); the Medical Research Council (M C_UU_00030/14; MR/T033371/1) and Cambridge Centre for Parkinson-plus; the Holt fellowship; and the NIHR Cambridge Biomedical Research Centre (NIHR203312: BRC-1215-20014). KJF is supported by funding for the Wellcome Centre for Human Neuroimaging (Ref: 205103/Z/16/Z) and a Canada-UK Artificial Intelligence Initiative (Ref: ES/T01279X/1). The views expressed are those of the author(s) and not necessarily those of the NIHR or the Department of Health and Social Care. For the purpose of open access, the authors have applied a CC BY public copyright licence to any Author Accepted Manuscript version arising from this submission.

## 8 Supplementary Figure

The graphics in this supplementary material are power spectral densities (PSD) and their associated predicated response by DCM at baseline and after two weeks in the eyes open condition, for the same subjects, at the four default mode network sources. Note that differences observed between baseline and re-test data two weeks later are not attributable to the progression of the disease in such a short time but may be related to other factors e.g. plasticity, psychological effects, differential fatigue, measurement noise or movement. Predicted simulated responses for the four-node default model network DCMs suggest that the neuronal model replicates most regions’ PSDs well. Note that DCM inversion gives weights or uncertainty to each frequency bin of PSDs, where intuitively low-frequency contents in observed signals may be considered less critical to essential brain rhythms, e.g., alpha, beta, etc. In addition, where a region’s activity (e.g. PCC with maximum alpha peak of 0.05 (*μv*^2)/*Hz*) is much smaller compared to another region (e.g. LAG with maximum PSD of 2 or 3 (*μv*^2)/*Hz*), the overall model likelihood and estimated signal-to-noise ratio (precision of the difference between model output and data) may be less sensitive.

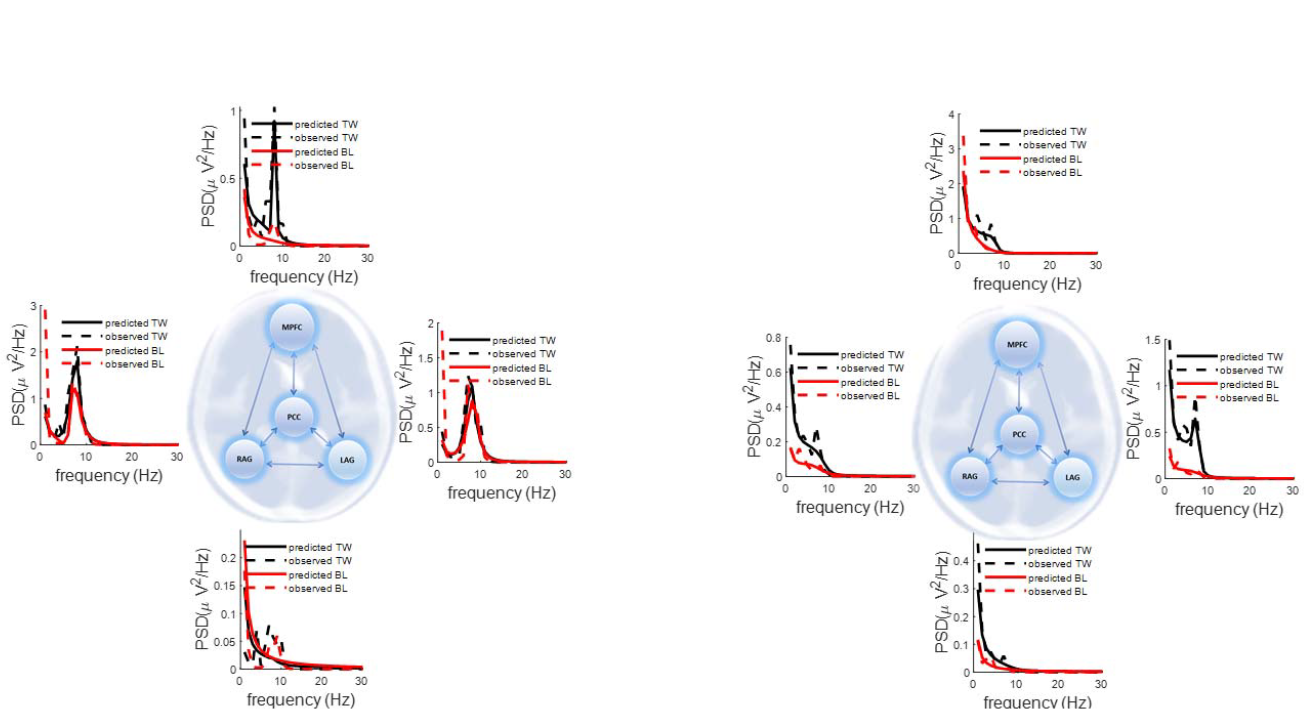
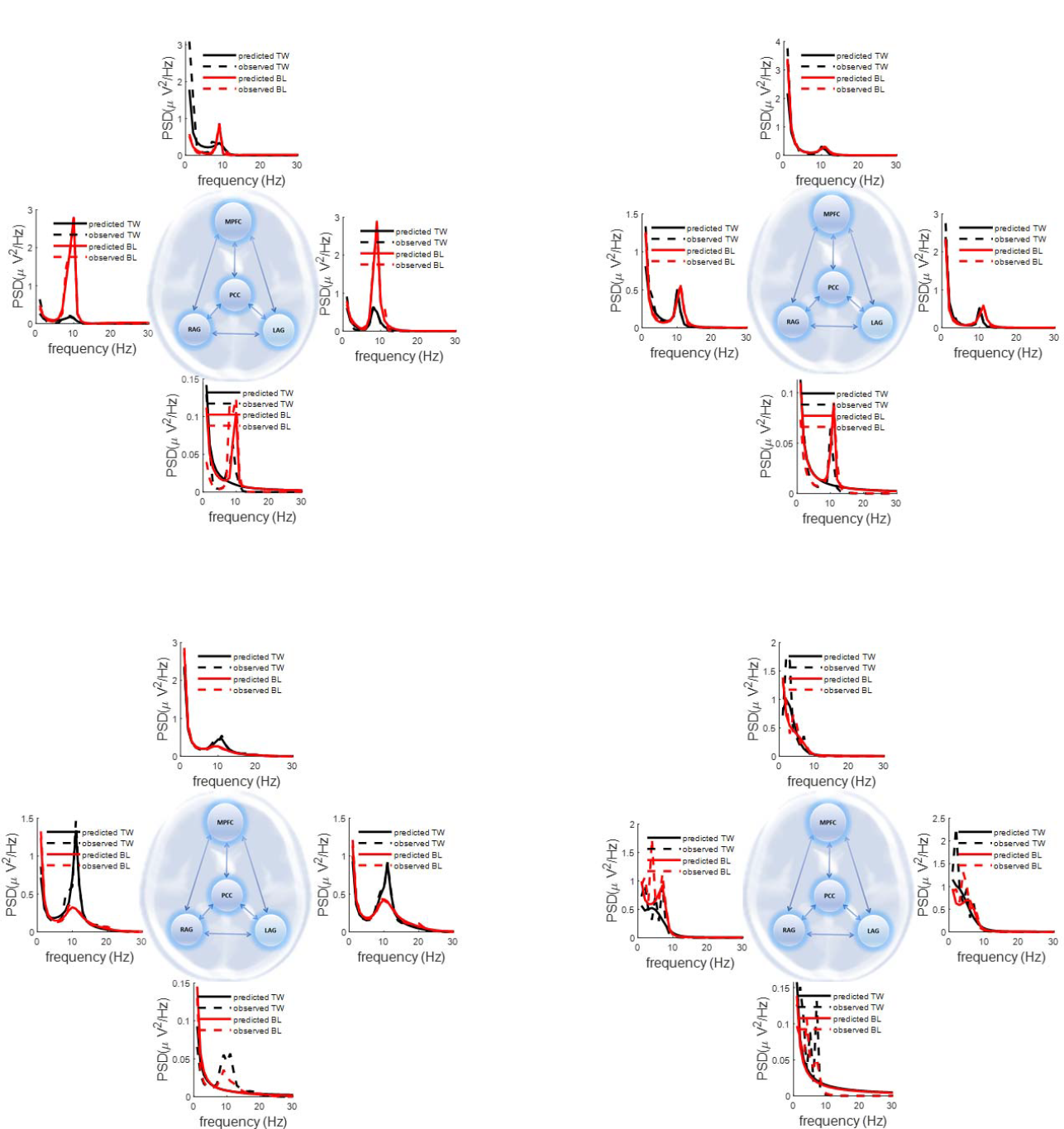
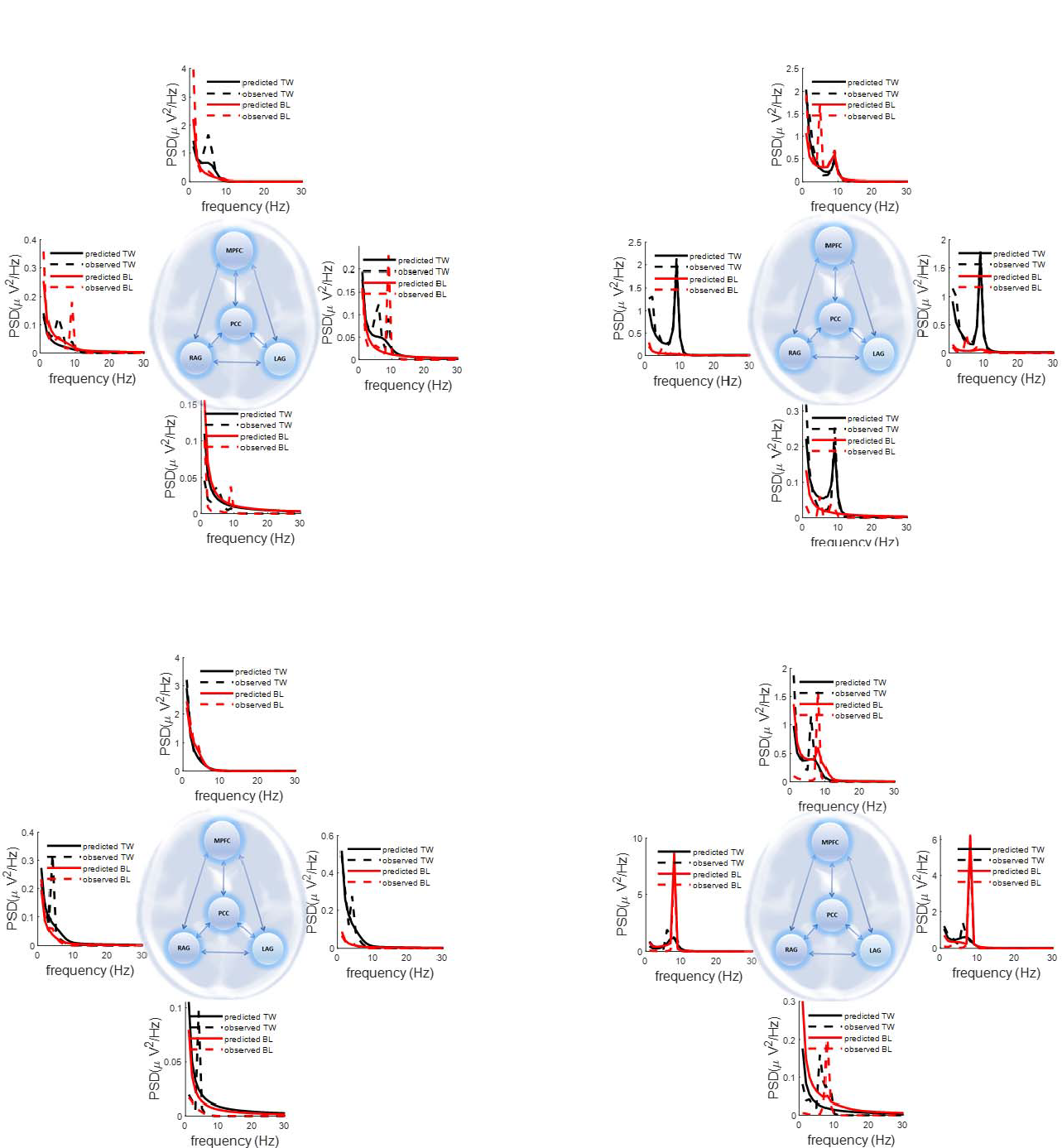
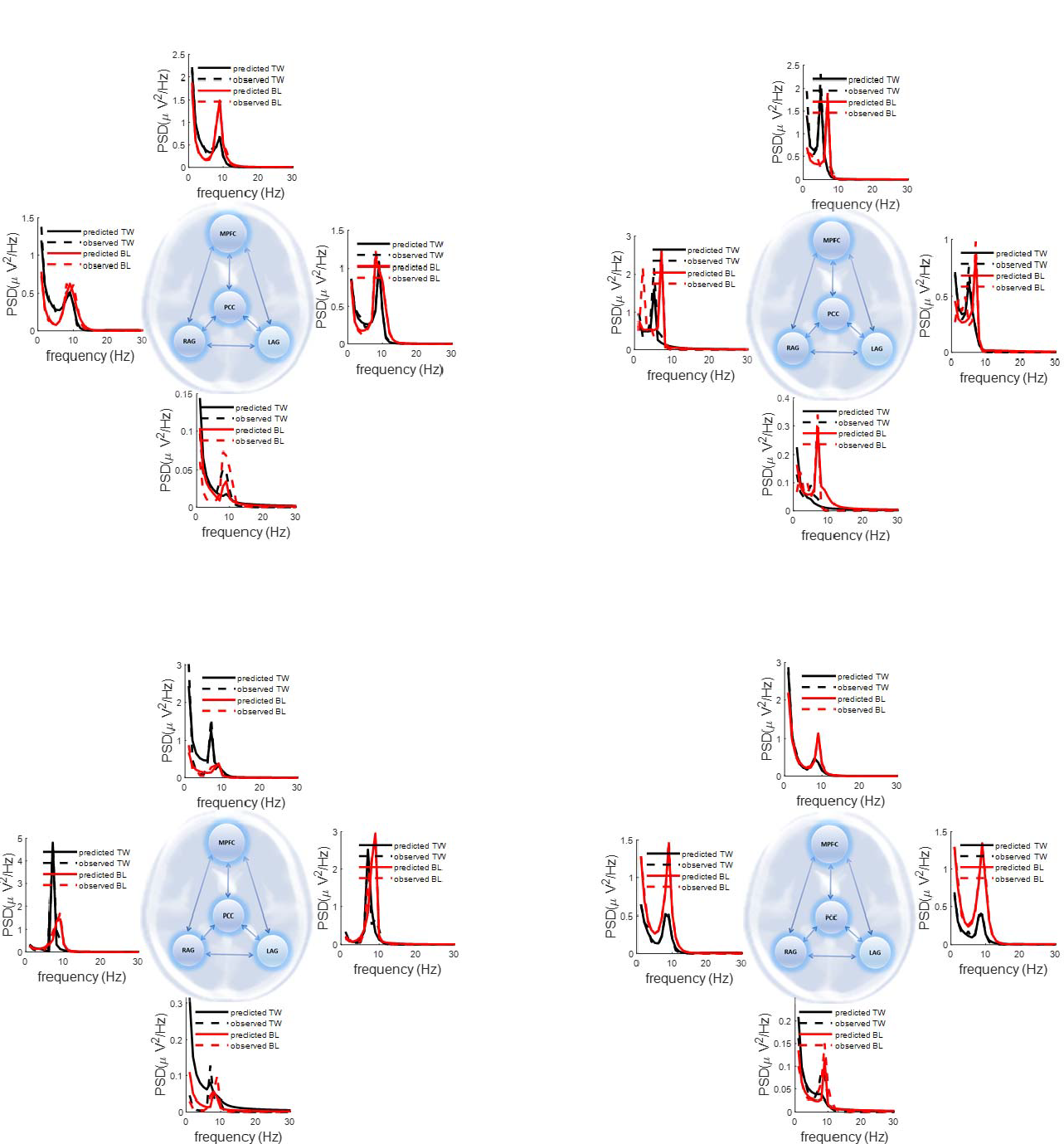

## Notes

### Competing Interest Statement

The authors have declared no competing interest.

## References

Adams, N. E., Hughes, L. E., Rouse, M. A., Phillips, H. N., Shaw, A. D., Murley, A. G., Cope, T. E., Bevan-Jones, W. R., Passamonti, L. & Street, D. 2021a. GABAergic cortical network physiology in frontotemporal lobar degeneration. Brain, 144, 2135–2145.

Adams, N. E., Jafarian, A., Perry, A., Rouse, M., Shaw, A. D., Murley, A. G., Cope, T. E., Bevan-Jones, W. R., Passamonti, L. & Street, D. 2022. Neurophysiological consequences of synapse loss in progressive supranuclear palsy. medRxiv.

Adams, R., Pinotsis, D., Tsirlis, K., Unruh, L., Mahajan, A., Horas, A., Convertino, L., Summerfelt, A., Sampath, H. & Du, X. 2021b. Computational modelling of EEG and fMRI paradigms indicates a consistent loss of pyramidal cell synaptic gain in schizophrenia. Biological Psychiatry.

Bartko, J. J. 1966. The intraclass correlation coefficient as a measure of reliability. Psychological reports, 19, 3–11.

Basar, E., Flohr, H., Haken, H. & Mandell, A. 2012. Synergetics of the Brain: Proceedings of the International Symposium on Synergetics at Schloß Elmau, Bavaria, May 2–7, 1983, Springer Science & Business Media.

Box, G. & Tiao, G. 1973. Bayesian Inference in Statistical Analyses Addison-Wesley-5 Longman. Reading, Massachusetts.

Fisher, R. A. 1992. Statistical methods for research workers. Breakthroughs in statistics. Springer.

FrÄSsle, S. & Stephan, K. E. 2022. Test-retest reliability of regression dynamic causal modeling. Network Neuroscience, 6, 135–160.

Friston, K., Mattout, J., Trujillo-Barreto, N., Ashburner, J. & Penny, W. 2007. Variational free energy and the Laplace approximation. Neuroimage, 34, 220–234.

Friston, K. & Penny, W. 2011. Post hoc Bayesian model selection. Neuroimage, 56, 2089–2099.

Friston, K., Zeidman, P. & Litvak, V. 2015. Empirical Bayes for DCM: a group inversion scheme. Frontiers in systems neuroscience, 9, 164.

Friston, K. J. 2011. Functional and effective connectivity: a review. Brain connectivity, 1, 13–36.

Friston, K. J., Bastos, A., Litvak, V., Stephan, K. E., Fries, P. & Moran, R. J. 2012. DCM for complex-valued data: cross-spectra, coherence and phase-delays. Neuroimage, 59, 439–455.

Friston, K. J., Flandin, G. & Razi, A. 2022. Dynamic causal modelling of COVID-19 and its mitigations. Scientific reports, 12, 12419.

Friston, K. J., Harrison, L. & Penny, W. 2003. Dynamic causal modelling. Neuroimage, 19, 1273–1302.

Friston, K. J., Li, B., Daunizeau, J. & Stephan, K. E. 2011. Network discovery with DCM. Neuroimage, 56, 1202–1221.

Friston, K. J., Litvak, V., Oswal, A., Razi, A., Stephan, K. E., Van Wijk, B. C., Ziegler, G. & Zeidman, P. 2016. Bayesian model reduction and empirical Bayes for group (DCM) studies. Neuroimage, 128, 413–431.

Friston, K. J., Preller, K. H., Mathys, C., Cagnan, H., Heinzle, J., Razi, A. & Zeidman, P. 2019. Dynamic causal modelling revisited. Neuroimage, 199, 730–744.

Friston, K. J., Trujillo-Barreto, N. & Daunizeau, J. 2008. DEM: a variational treatment of dynamic systems. Neuroimage, 41, 849–885.

Gilbert, J. R., Symmonds, M., Hanna, M. G., Dolan, R. J., Friston, K. J. & Moran, R. J. 2016. Profiling neuronal ion channelopathies with non-invasive brain imaging and dynamic causal models: case studies of single gene mutations. Neuroimage, 124, 43–53.

Greicius, M. D., Srivastava, G., Reiss, A. L. & Menon, V. 2004. Default-mode network activity distinguishes Alzheimer’s disease from healthy aging: evidence from functional MRI. Proceedings of the National Academy of Sciences, 101, 4637–4642.

Haken, H. 1977. Synergetics. Physics Bulletin, 28, 412.

Jafarian, A., Hughes, L., Adams, N. E., Lanskey, J., Naessens, M., Rouse, M. A., Murley, A. G., Friston, K. & Rowe, J. 2022. Neurochemistry-enriched dynamic causal models of magnetoencephalography, using magnetic resonance spectroscopy. bioRxiv.

Jafarian, A., Hughes, L. E., Adams, N. E., Lanskey, J., Naessens, M., Rouse, M. A., Murley, A. G., Friston, K. J. & Rowe, J. B. 2023. Neurochemistry-enriched dynamic causal models of magnetoencephalography, using magnetic resonance spectroscopy. NeuroImage, 120193.

Jafarian, A., Zeidman, P., Litvak, V. & Friston, K. 2019. Structure learning in coupled dynamical systems and dynamic causal modelling. Philosophical Transactions of the Royal Society A, 377, 20190048.

Jafarian, A., Zeidman, P., Wykes, R. C., Walker, M. & Friston, K. J. 2021. Adiabatic dynamic causal modelling. NeuroImage, 238, 118243.

Jelenkowska, T. H. 1998. Bayesian estimation of the intraclass correlation coefficients in the mixed linear model. Applications of Mathematics, 43, 103–110.

Jelenkowska, T. H. 1999. Bayesian estimation of the intraclass correlation coefficients in the multivariate one-way model. Listy Biometryczne-Biometrical Letters, 36, 61–67.

Kass, R. E. & Raftery, A. E. 1995. Bayes factors. Journal of the american statistical association, 90, 773–795.

Lanskey, J. H., Kocagoncu, E., Quinn, A. J., Cheng, Y.-J., Karadag, M., Pitt, J., Lowe, S., Perkinton, M., Raymont, V. & Singh, K. D. 2022. New Therapeutics in Alzheimer’s Disease longitudinal cohort study (NTAD): study protocol. BMJ open, 12, e055135.

Litvak, V., Garrido, M., Zeidman, P. & Friston, K. 2015. Empirical Bayes for group (DCM) studies: a reproducibility study. Frontiers in human neuroscience, 9, 670.

Litvak, V., Jafarian, A., Zeidman, P., Tibon, R., Henson, R. N. & Friston, K. There’s no such thing as a ‘true’model: the challenge of assessing face validity. 2019 IEEE International Conference on Systems, Man and Cybernetics (SMC), 2019. IEEE, 4403–4408.

Litvak, V., Mattout, J., Kiebel, S., Phillips, C., Henson, R., Kilner, J., Barnes, G., Oostenveld, R., Daunizeau, J. & Flandin, G. 2011. EEG and MEG data analysis in SPM8. Computational intelligence and neuroscience, 2011.

Lorenzi, M., Beltramello, A., Mercuri, N. B., Canu, E., Zoccatelli, G., Pizzini, F. B., Alessandrini, F., Cotelli, M., Rosini, S. & Costardi, D. 2011. Effect of memantine on resting state default mode network activity in Alzheimer’s disease. Drugs & aging, 28, 205–217.

Moran, R. J., Kiebel, S. J., Stephan, K. E., Reilly, R., Daunizeau, J. & Friston, K. J. 2007. A neural mass model of spectral responses in electrophysiology. NeuroImage, 37, 706–720.

Moran, R. J., Pinotsis, D. A. & Friston, K. J. 2013. Neural masses and fields in dynamic causal modeling. Frontiers in computational neuroscience, 7, 57.

Moran, R. J., Stephan, K. E., Dolan, R. J. & Friston, K. J. 2011. Consistent spectral predictors for dynamic causal models of steady-state responses. Neuroimage, 55, 1694–1708.

Mulder, J. & Fox, J.-P. 2019. Bayes factor testing of multiple intraclass correlations. Bayesian Analysis, 14, 521–552.

Penny, W. D. 2012. Comparing dynamic causal models using AIC, BIC and free energy. Neuroimage, 59, 319–330.

Prinz, A. A., Bucher, D. & Marder, E. 2004. Similar network activity from disparate circuit parameters. Nature neuroscience, 7, 1345–1352.

Rowe, J. B. 2010. Connectivity analysis is essential to understand neurological disorders. Frontiers in Systems Neuroscience, 4, 144.

Rowe, J. B., Hughes, L. E., Barker, R. A. & Owen, A. M. 2010. Dynamic causal modelling of effective connectivity from fMRI: are results reproducible and sensitive to Parkinson’s disease and its treatment? Neuroimage, 52, 1015–1026.

Schuyler, B., Ollinger, J. M., Oakes, T. R., Johnstone, T. & Davidson, R. J. 2010. Dynamic causal modeling applied to fMRI data shows high reliability. Neuroimage, 49, 603–611.

Shaw, A., Moran, R. J., Muthukumaraswamy, S. D., Brealy, J., Linden, D., Friston, K. J. & Singh, K. D. 2017. Neurophysiologically-informed markers of individual variability and pharmacological manipulation of human cortical gamma. Neuroimage, 161, 19–31.

Shaw, A. D., Hughes, L. E., Moran, R., Coyle-Gilchrist, I., Rittman, T. & Rowe, J. B. 2021. In vivo assay of cortical microcircuitry in frontotemporal dementia: A platform for experimental medicine studies. Cerebral cortex, 31, 1837–1847.

Stephan, K. E., Tittgemeyer, M., KnÖSche, T. R., Moran, R. J. & Friston, K. J. 2009. Tractography-based priors for dynamic causal models. Neuroimage, 47, 1628–1638.

Stephan, K. E., Weiskopf, N., Drysdale, P. M., Robinson, P. A. & Friston, K. J. 2007. Comparing hemodynamic models with DCM. Neuroimage, 38, 387–401.

Wang, M. & Sun, X. 2014. Bayes factor consistency for one-way random effects model. Communications in Statistics-Theory and Methods, 43, 5072–5090.

Zeidman, P., Jafarian, A., Seghier, M. L., Litvak, V., Cagnan, H., Price, C. J. & Friston, K. J. 2019. A guide to group effective connectivity analysis, part 2: Second level analysis with PEB. NeuroImage, 200, 12–25.

